# Press xenobiotic disturbance favors deterministic assembly with a shift in function and structure of bacterial communities in sludge bioreactors

**DOI:** 10.1101/2020.10.15.341966

**Authors:** Ezequiel Santillan, Hari Seshan, Stefan Wuertz

**Affiliations:** Singapore Centre for Environmental Life Sciences Engineering, Nanyang Technological University, 637551, Singapore; Department of Civil and Environmental Engineering, University of California, Davis, CA 95616, U.S.A; School of Civil and Environmental Engineering, Nanyang Technological University, 639798, Singapore

**Keywords:** diversity, disturbance, community structure, stochastic assembly, deterministic assembly

## Abstract

Disturbance is thought to affect community assembly mechanisms, which in turn shape community structure and the overall function of the ecosystem. Here, we tested the effect of a continuous (press) xenobiotic disturbance on the function, structure, and assembly of bacterial communities within a wastewater treatment system. Two sets of four-liter sequencing batch reactors were operated in triplicate with and without the addition of 3-chloroaniline for a period of 132 days, following 58 days of acclimation after inoculation with sludge from a full-scale treatment plant. Temporal dynamics of bacterial community structure were derived from 16S rRNA gene amplicon sequencing. Community function, structure and assembly differed between press disturbed and undisturbed reactors. Temporal partitioning of assembly mechanisms via phylogenetic and non-phylogenetic null modelling analysis revealed that deterministic assembly prevailed for disturbed bioreactors, while the role of stochastic assembly was stronger for undisturbed reactors. Our findings are relevant because research spanning various disturbance types, environments and spatiotemporal scales is needed for a comprehensive understanding of the effects of press disturbances on assembly mechanisms, structure, and function of microbial communities.

**Graphical abstract:** 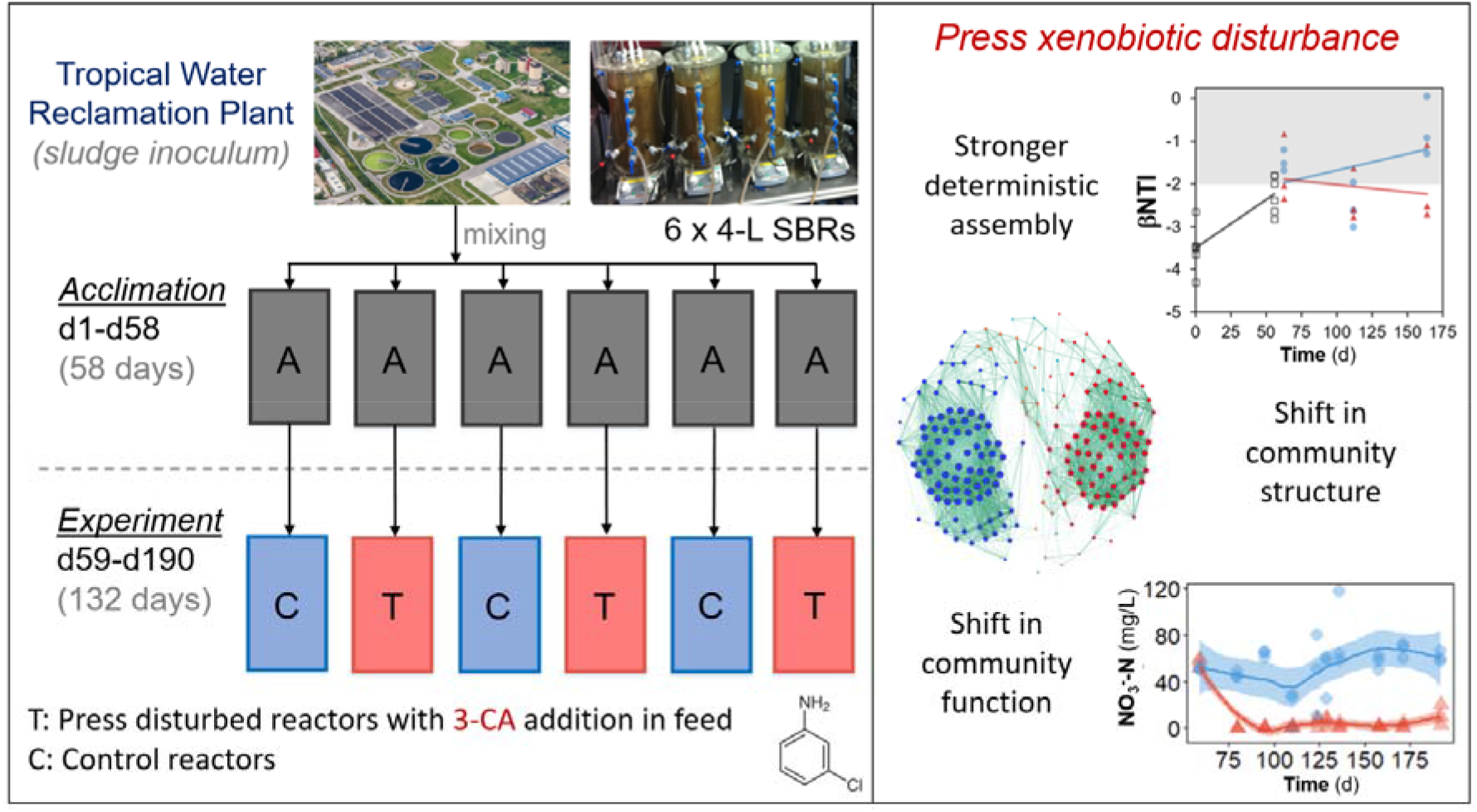

## Introduction

Sludge bioreactors for wastewater treatment are good model systems for microbial ecology studies^1^. These engineered systems involve measurable functions such as the removal of carbon and ammonia that are relevant in practice^2^ and entail complex microbial communities in a controlled environment^3^. In nature, microbial communities drive all biogeochemical cycles globally while providing ecosystem functions necessary for all other life forms to exist^4^. The structure of these communities, which can be described in terms of α- and β-diversity metrics, is believed to shape the ecosystem function they provide^5^. Predicting and managing the functions of microbial communities based on a fundamental understanding of their relationship with community structure remains a key challenge for complex microbial systems.

Disturbance is thought to have direct impacts on ecosystems by shifting community structure and function^6^. Chloroanilines are intermediate breakdown products from the use of herbicides and pesticides in agriculture^7^, which can be found in rubber, dye, polymer and pharmaceutical industrial wastewater treatment plants^8,9^, but also in soil and natural aqueous environments^10^. These xenobiotic compounds are known to hamper both carbon and nitrogen removal in bioreactors for wastewater treatment^11^, and can represent a disturbance in sludge bioreactor systems. Press disturbances, which inflict a long-term continuous alteration of taxa abundances by changing their environment^12^, are relevant for both microbial ecology and environmental biotechnology as they can lead systems to alternative stable states with different community function and structure^13^.

Community assembly processes are fundamentally related to ecosystem function, as these are thought to shape community structure^14^. These processes can be either deterministic or stochastic, often acting in combination to shape patterns of community assembly^15–17^. In ecology, the contribution of assembly mechanisms is usually quantified via null model analyses^18^. Although disturbance is thought to be a main driver of these underlying community assembly mechanisms^19^, a predictive understanding of its effects remains elusive^20^. Disturbance can favour stochastic assembly mechanisms that could lead communities to deviating states of function and structure^21,22^; hence, assessment of its effects demands replicated study designs^23,24^. Further, while several studies have reported patterns of community assembly in engineered microbial systems^25–28^, relatively few have addressed the effects of disturbance on community assembly, structure and function of such systems, particularly for bioreactors treating wastewater.

The objective of this work was to test the effect of a press disturbance on community assembly mechanisms by introducing a xenobiotic compound, 3-chloroaniline (3-CA), in a replicated set of activated sludge bioreactors with a working volume of four liters, representing a mesocosm scale. Based on our findings in a prior reactor study at a microcosm scale^21^, we expected to see a stronger deterministic effect at the disturbed level. Samples were analysed using 16S rRNA gene amplicon sequencing and effluent chemical characterization. Patterns of α- and β-diversity were employed to assess temporal dynamics of community structure. Assembly mechanisms were quantified via two different mathematical null models, one that assessed the phylogenetic turnover for each bioreactor, and another that evaluated the effective bacterial turnover expressed as a proportion of total bacterial diversity^29^ across all replicate reactors.

## Materials and Methods

### Experimental design

Six sequencing batch bioreactors (SBRs) were operated in parallel and fed synthetic wastewater for an acclimation period of 58 d. The inoculum was taken from the aeration tanks of a full-scale wastewater treatment plant (WWTP) in Singapore. On day 0 of acclimation, the freshly sourced inoculum sludge was mixed well in the laboratory and distributed to all six reactors (2 L each). Each was topped up with 2 L of synthetic wastewater, to a working volume of 4 L per reactor. The reactors were run under identical conditions with 12-h cycles as follows: 20 min of feeding, 180 min of anoxic mixing, 440 min of aeration and mixing (dissolved oxygen, DO, maintained at 1-2 mg/L using a feedback loop where aeration would commence at 1 L air/min when probes measured DO as below 1 mg/L and aeration would stop when the DO reading was above 2 mg/L), 50 min of settling and 30 min of effluent (supernatant) discharge. Two liters of effluent were discharged at the end of every cycle and replaced with 2 L of synthetic wastewater at the beginning of the next 12-h cycle, resulting in a hydraulic retention time (HRT) of 48 h. The mixed liquor temperature was maintained at 30°C using water jackets around the reactors and a re-circulating water heater. Solids were removed regularly from the mixed liquor to maintain a solids retention time (SRT) of about 30 d in each reactor.

The synthetic wastewater fed to all six reactors during the acclimation period was adapted from Hesselmann *et al.*^30^ and contained the following (mg/L in in each reactor after feeding): sodium acetate (112.5), dextrose (45), yeast extract (67.5), soy peptone (60), meat peptone (60), casein peptone (90), urea (15), ammonium bicarbonate (90), ammonium chloride (169), disodium hydrogen phosphate (720), potassium dihydrogen phosphate (130), calcium chloride dihydrate (10.5) and magnesium sulphate heptahydrate (112.5). The medium also contained 2 mL of the unaltered trace element stock^30^ per liter of medium. The first six components contributed to the COD, amounting to about 500 mg/L in the reactor, the next three (ammonium-based) components contributed to the total loading of about 70 mg N/ L, and the phosphates were used to buffer the medium and maintain a pH of around 7.5 to facilitate the nitrification process.

After the 58-d acclimation period, the sustained 3-CA input experiment was started and continued for 132 d. At the start of this experiment, three of the acclimated reactors were randomly assigned to the treatment group (press disturbed with 3-CA) and the other three were assigned to the control group (no 3-CA addition, or undisturbed). The cycle conditions and other parameters were kept the same among all six reactors and were identical to the conditions in the acclimation phase. The 3-CA inputs to the treatment reactors started on day 59. The medium used to feed the treatment reactors was slightly altered: the organic constituents (sodium acetate, dextrose, yeast extract, soy peptone, meat peptone and casein peptone) were scaled down by 20%, resulting in a total COD of 400 mg/L from these constituents. The remaining 20% of the COD was fed in the form of 3-CA, resulting in a mixed liquor 3-CA concentration of about 70 mg/L, an environmentally relevant level for WWTPs^11^. The three control reactors continued to receive the same 3-CA-free medium that was used during acclimation.

To assess the changes in community composition, samples of sludge were taken from all six reactors at five distinct time points, representing five different periods in the progression of the experiment. These five time points were day 0 (the day the reactors were inoculated for acclimation), day 56 (shortly before 3-CA addition to treatment reactors was commenced), day 63 (shortly after 3- CA addition to treatment reactors was commenced), day 112 (after process performance had stabilised) and day 164 (toward the end of the experiment). On each of these sampling days, sludge was collected directly from the mixed liquor at the beginning of a given anoxic phase, aliquoted into 1 mL in cryogenic tubes, flash-frozen in liquid nitrogen and stored at −80°C for subsequent molecular analysis via 16S rRNA gene amplicon sequencing and terminal restriction fragment length polymorphism (T-RFLP) analysis. The T-RFLP results are discussed elsewhere^31^.

### Water chemical analysis

Water quality parameters were measured using standard nutrient analysis techniques in accordance with standard methods^32^ and targeted COD (standard methods 5220 D), nitrogen species (ammonium, nitrite and nitrate ions) using ion chromatography (standard methods 4500-NH3 for ammonium; 4110 B for nitrate and nitrite). Nitrogen species were also measured using spectrophotometric tests (Hach) to complement ion chromatography results. 3-CA was measured on a Shimadzu Prominence High Pressure Liquid Chromatography system (Shimadzu) equipped with a UV-VIS PDA detector using an Ascentis C18 5-μm column (Sigma-Aldrich). An isocratic 50:50 Water:Methanol solvent was used at a flow of 0.3 mL/min, and 3-CA peaks were identified and measured at 199, 237 and 286 nm.

### 16S rRNA gene amplicon sequencing and reads processing

Genomic DNA was extracted from samples collected from bioreactors on the five days identified above (500 μL of sludge per sample) using the FastDNA Spin Kit for Soil (MP Biomedicals) with alterations to the manufacturer’s protocol to increase DNA yield as detailed elsewhere^21^. Extracted DNA was amplified in triplicate 50-μL PCR reactions using primer set 530f/U1053r, which targets the V3~V4-V5 variable regions of the bacterial 16S rRNA gene^33^. Each 50-μL PCR reaction contained 25 μL of ImmoMix reagent (Meridian Bioscience) and about 200 ng of DNA extract and nuclease-free water. The PCR program included an initial denaturation step at 95° C for 10 min, followed by 30 cycles of denaturation (95°C, 1 min), annealing (58°C, 30 s) and extension (72°C, 1 min). A final extension was carried out at 72° C for 7 min. The triplicate PCR amplicons from each sample-primer set combination were then pooled and purified using the QIAquick PCR purification kit (Qiagen), with a slightly altered protocol in which the final elution was performed on nuclease-free water pre-heated to 55°C and incubated for 5 min at 55° C before the final centrifugation. This modification was found to increase the final DNA yield. The purified amplicons were then inspected for quality using agarose gels and quantified using a Qubit 3.0 fluorometer (ThermoFisher Scientific).

The libraries were sequenced in-house at SCELSE on a Illumina MiSeq (v.3) with 20% PhiX spike-in, at 300 bp paired-end read-length. Sequenced sample libraries were processed with the *dada2* (v.1.3.3) R-package^34^, allowing inference of amplicon sequence variants (ASVs)^35^. Illumina adaptors and PCR primers were trimmed prior to quality filtering. Sequences were truncated after 280 and 255 nucleotides for forward and reverse reads, respectively. After truncation, reads with expected error rates higher than 3 and 5 for forward and reverse reads, respectively, were removed. Since paired-end reads could not be merged with a minimum overlap of 20 bp, only forward reads were kept for further analysis. Chimeric sequences (0.3% on average) were identified and removed. For a total of 30 samples, an average of 48,659 reads were kept per sample after processing, representing 61% of the average forward input reads. Taxonomy was assigned using the SILVA database (v.132)^36^. Samples were rarefied to the lowest number of reads (18,353) in a sample after processing (Fig. S1).

### Bacterial community analysis and statistics

All reported p-values for statistical tests in this study were corrected for multiple comparisons using a False Discovery Rate (FDR) of 5%^37^. Community structure was assessed by a combination of principal coordinate analysis (PCoA) ordination of weighed Unifrac dissimilarity matrixes, constructed from Hellinger transformed abundance data using the *phyloseq^38^* package in R, and multivariate tests of permutational analysis of variance (PERMANOVA) and permutational analysis of dispersion (PERMDISP) on Bray-Curtis dissimilarity matrixes, constructed from square-root transformed abundance data using PRIMER (v.7)^39^. Hill diversity indices^40^ were used to quantify α- diversity as described elsewhere^21^. Local polynomial regression fitting was applied using the *loess* function from the *ggplot2* package in R^41^, including 95% confidence intervals. Welch’s t-test (twotailed) was used for univariate testing. Heat maps for bacterial genera relative abundances were constructed using the *ampvis2* package in R^42^. Network analysis was performed on data from day 164 with Gephi (v.0.9.2) using Spearman’s correlation matrixes for the top 200 ASVs. Input correlation matrixes were generated using the *Hmisc* and *corrplot* R packages. Non-significant correlations (ρ < 0.6) were filtered out. Node clusters were defined by modularity class calculated using the Louvain method, and were coloured through identification of representative nodes at undisturbed and disturbed levels. Layout was adjusted using the Fruchterman Reingold method with default parameters. Node size was adjusted by degree, while edge thickness was adjusted by correlation strength.

### Null model analyses on diversity

The effect of underlying assembly mechanisms was first assessed using a phylogenetic-based null modelling approach. The model uses the β-mean nearest taxon distance (βMNTD)^43^ which quantifies the phylogenetic distance between each ASV in one community, as a measure of the clustering of closely related ASVs. Phylogenetic relatedness of ASVs was characterized by multiple-alignment of ASV sequences using *decipher* R-package^44^. The phylogenetic tree was then constructed and a GTR+G+I maximum likelihood tree was fitted using the *phangorn* R-package^45^. To quantify the degree to which βMNTD deviates from a null model expectation, ASVs and abundances were shuffled across the tips of the phylogenetic tree. After shuffling, βMNTD was recalculated to obtain a null value, and repeating the shuffling 1,000 times provided a null distribution. The difference between observed βMNTD and the mean of the null distribution was measured in units of standard deviation, which is referred to as the β-nearest taxon index (βNTI)^46^. A value of |βNTI| > 2 indicates that the observed turnover between a pair of communities is significantly deterministic, while |βNTI| < 2 suggests stochastic assembly^16^. This analysis was done using the *phylocom* R-package^47^.

We further assessed assembly mechanisms by using an alternative null modelling methodology, which assumes that species interactions are not important for community assembly^18^ and quantifies the effective bacterial turnover expressed as a proportion of total bacterial diversity. It was developed for woody plants and recently applied to sludge ‘and groundwater microbial communities. The model defines β-diversity as the β-partition 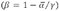. To adapt it to handle microbial community data, we considered ‘species’ in the model as ASVs, while each individual count was one read within the corresponding dataset. The model randomizes the location of each individual within the independent replicate reactors for each of the control and treatment levels, while maintaining the total quantity of individuals per reactor, the relative abundance of each ‘species’*(i.e.,* ASVs), and the γ-diversity. This way it takes into account both composition and relative abundances. We applied the model across different time points of the experiment. Control and treatment levels had three replicate reactors each, thus acclimation phase data were assigned to the same two groups of three reactors each for model estimations. Each step of the null model calculated expected mean α-diversities per treatment level and then estimated an expected β-partition. After 10,000 repetitions, the means of the distribution of random β-partitions 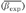 for each treatment level were calculated. The relative contribution of deterministic assembly mechanisms was then quantified using the stochastic intensity (SI) metric, which measured the deviation of the observed β-diversity compared to that expected by chance as follows: 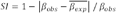. SI is the complement of the deterministic strength (DS) metric 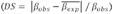, defined previously^17,21^. Higher values of SI indicated a lower deviation of the observed β-diversity from the null β-diversity expectation, thus suggesting a stronger effect of stochastic-based mechanisms. Contrarily, lower SI values indicated a bigger difference between observed and null β-diversities, suggesting a more important role of deterministic mechanisms of assembly.

## Results

### Dynamics of process performance

There was a clear distinction in process performance (*i.e.* ecosystem function) between control and treatment levels, with disturbed reactors displaying higher COD, 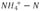 and 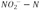 concentrations in the effluent due to inhibition of nitrification (Fig. 1A-C). Nitrate was found in the effluent of control reactors, which together with low levels of residual ammonia and nitrite indicated complete nitrification (Fig. 1D). During the course of the experiment the microbial communities in the three treatment reactors acquired the ability to metabolize 3-CA and perform partial nitrification (Fig. 1E). A detailed analysis of changes in process performance under 3-CA disturbance in this study is reported elsewhere^31^.

**Figure 1.**
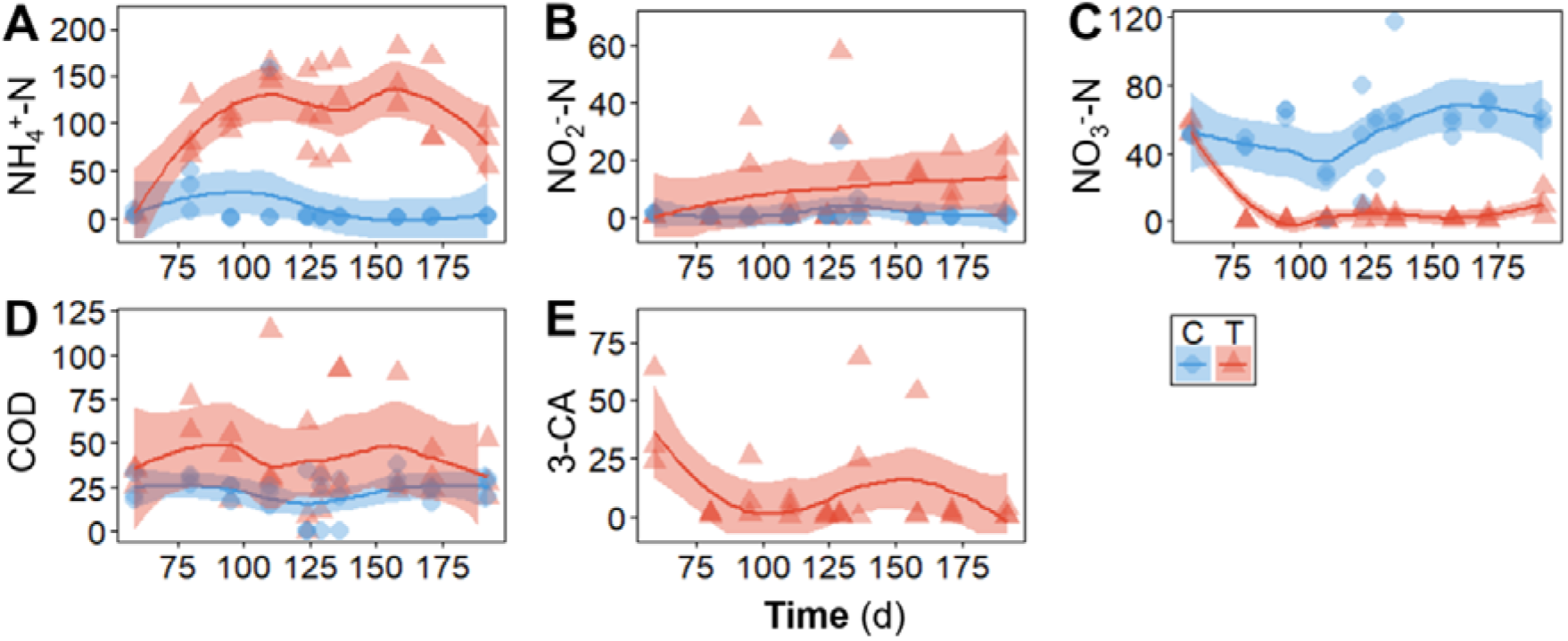
Temporal bioreactor effluent concentrations (mg/L) at the end of a reactor cycle, after the acclimation period (from d59 onwards). (**A**) Ammonium, (**B**) nitrite, and (**C**) nitrate as nitrogen, (**D**) organic carbon as soluble chemical oxygen demand (COD) and (**E**) 3-chloroaniline. Each point represents a different reactor for a given day. Reactor type: C, control (blue, n = 3); T, treatment (red, n = 3). Lines display polynomial regression fitting, while shaded areas represent 95% confidence intervals. No 3-CA was fed to control reactors, thus no blue points displayed in panel **E**.

### Dynamics of bacterial community structure

Throughout the acclimation phase there were significant changes in bacterial community structure, both in terms of α-diversity (Fig. 2) (P-_Welch’s t-test_< 0.009) and β-diversity (Fig. 3) (P-_PERMANOVA_ < 0.0001), from the initial wastewater treatment plant inoculum (day 0) all the way to the acclimated sludge on day 58. α-diversity after the acclimation period varied over time, but did not differ significantly between control and treatment reactors (P-_Welch’s t-test_ > 0.10), although control reactors displayed higher α-diversity only towards the end of the study (Fig. 2A-C). However, temporal patterns of β-diversity showed bacterial communities clustering separately in control and treatment reactors (Fig. 3). Such differentiation was statistically significant from day 112 onwards (P-_PERMANOVA_ < 0.016), with no effects of heteroscedasticity (P-_PERMDISP_ > 0.41, Table S1). Bacterial succession was also evident from the analysis of relative abundances of specific taxa, which were different at each stage of the study (Fig. 4A). Genera like *Propioniciclava, OLB8* and *Terrimonas* prevailed initially on day 0. After acclimation *Paracoccus*, *Kouleothrix* and *Alicycliphilus* dominated across control reactors, while *Ca.* Competibacter, *Dokdonella* and *Gemmatimonas* prevailed in treatment reactors. Some taxa like *Tetrasphaera* and *Microlunatus* increased in relative abundance for all reactors throughout the study. Correlation network analysis on day 164 also displayed separate clusters of bacterial ASVs based on node modularity (Fig. 4B).

**Figure 2.**
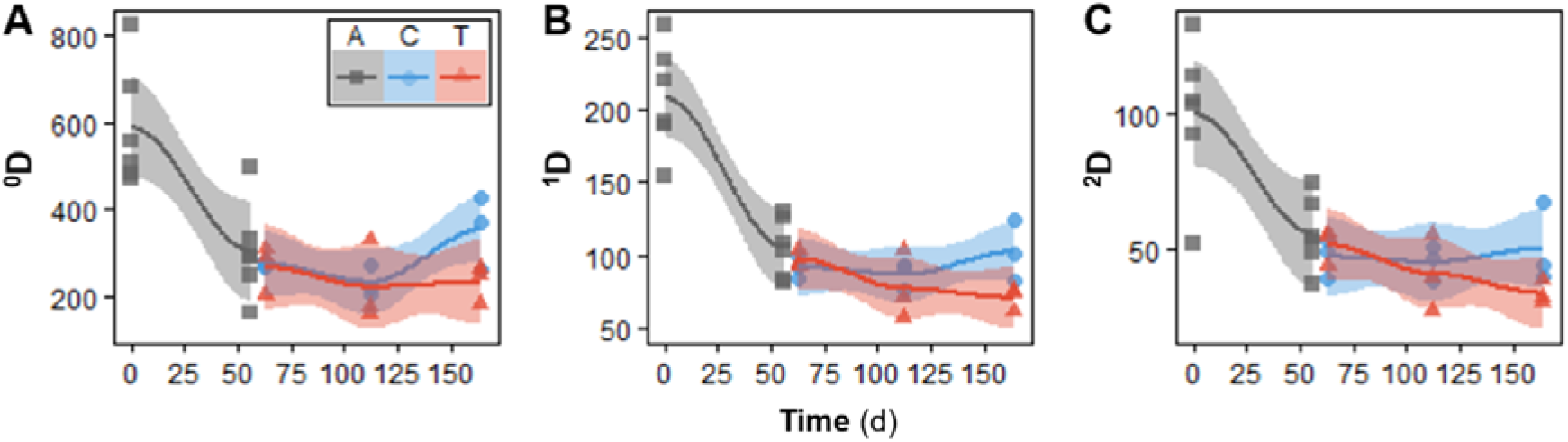
Temporal dynamics of true α-diversity Hill numbers for bacterial ASVs. Hill number orders: (**A**) ^0^D, (**B**) ^1^D, and (**C**) ^2^D. Each point represents a different reactor on a given day. Reactor type: A, acclimation (grey, n = 6); C, control (blue, n = 3); T, treatment (red, n = 3). Lines refer to polynomial regression fitting, while shaded areas represent 95% confidence intervals.

**Figure 3.**
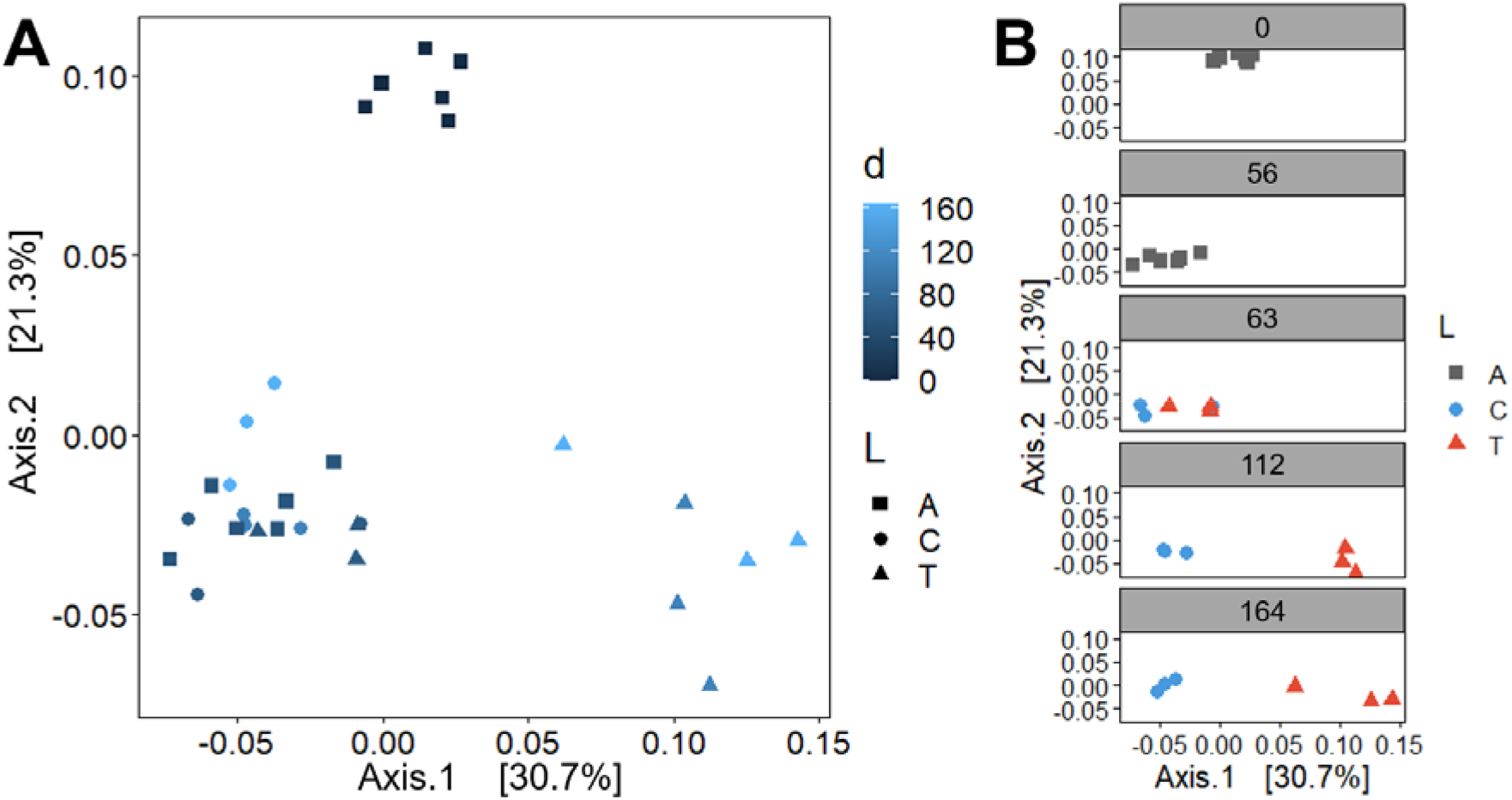
Temporal dynamics of community structure for bacterial ASVs evaluated through PCoA ordination (weighed Unifrac β-diversity). Panels: (**A**) all time points (time increases from dark to light blue) and (**B**) separate time points. Reactor type: A, acclimation (squares, n = 6); C, control (circles, n = 3); T, treatment (triangles, n = 3). Each point represents a different reactor for a given day.

**Figure 4.**
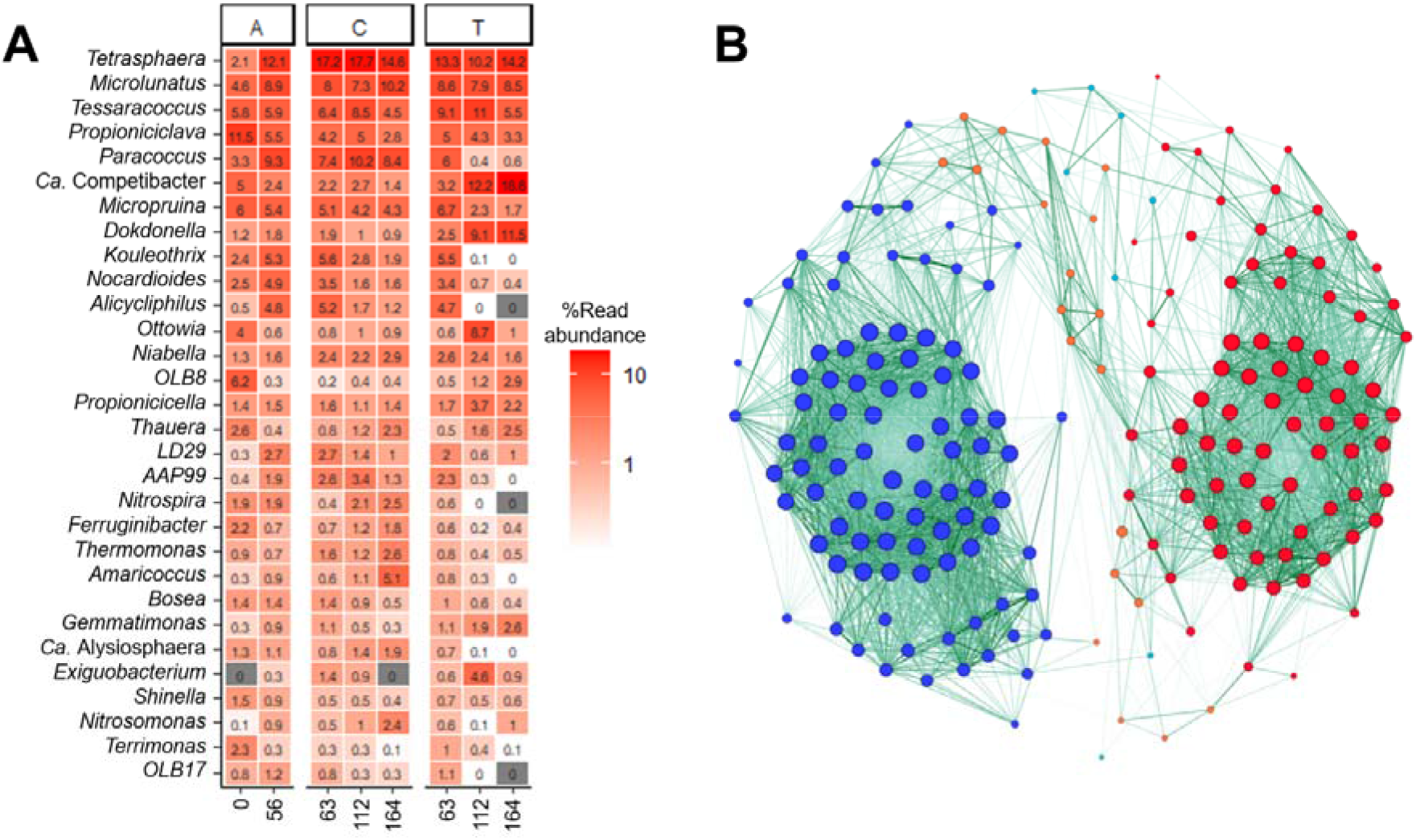
(**A**) Community structure dynamics for bacterial genera, assessed through 16S rRNA gene amplicon sequencing. The 30 most abundant genera are shown. Columns represent the average percentage read abundance among reactors for a given reactor type and day. Reactor type: A, acclimation (left, n = 6); C, control (middle, n = 3); T, treatment (right, n = 3). (**B**) Correlation network for the top 200 most abundant bacterial ASVs on d164. Clusters are coloured by modularity class, with blue nodes prevailing in control reactors (n = 3) and red nodes in treatment reactors (n = 3). Only significantly strong Spearman’s correlations (p ≥ 0.60) were employed. Edge thickness represents correlation strength and node size represents degree.

### Dynamics of bacterial community assembly mechanisms

Two different approaches were applied to quantify assembly mechanisms, based on the deviation of observed β-diversity from the expected β-diversity from null modelling. Phylogenetic turnover was first assessed using the β-nearest taxon index (βNTI). Treatment reactors had mostly mean values of βNTI < −2, indicating deterministic assembly (Fig. 5A). For control reactors, values of |βNTI| < 2 indicated a significant effect of stochastic assembly (Fig. 5A). The observed βNTI patterns were not driven by a particular reactor (Fig. S2). Concurrently, the effective bacterial turnover expressed as a proportion of total bacterial diversity was quantified via the stochastic intensity (SI) metric. Similarly, SI values trended higher in control reactors, suggesting a greater relative role of stochastic assembly mechanisms compared to treatment reactors (Fig. 5B).

**Fig. 5.**
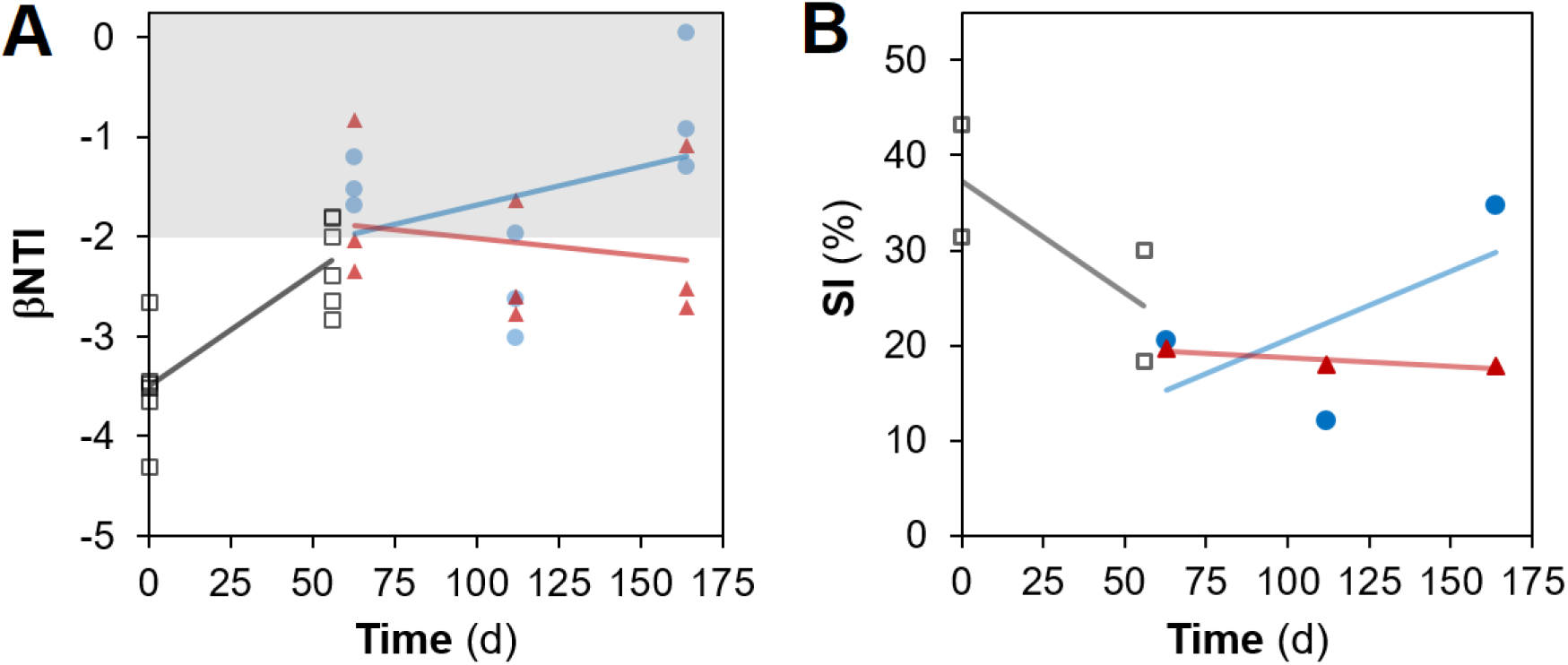
Temporal dynamics of community assembly for bacterial ASVs evaluated via (**A**) nearest taxon index (βNTI) and (**B**) stochastic intensity (SI), from null model analyses. Reactor type: A, acclimation (grey, n = 6); C, control (blue, n = 3); T, treatment (red, n = 3). Lines represent linear regression fitting. Shaded in grey in (**A**) is the zone where stochastic processes significantly dominate |βNTI| < 2. Each point in (**B**) involves three replicates for SI calculation.

## Discussion

For this study, we hypothesized that press disturbance by a xenobiotic would alter function, structure and assembly of sludge bioreactor bacterial communities. Chloroanilines are toxic substances that can easily diffuse within the natural environment and are difficult to remediate, posing an environmental problem for natural waters and soils where their concentrations are increasing^10^. This work used 3-CA as disturbance, a compound known to have a detrimental effect on relevant functions within wastewater bacterial communities^11,21^. Indeed, function (in terms of process performance) and structure (in terms of β-diversity) clearly differed between control and treatment reactors. Disturbance impaired nitrification in treatment reactors (Fig. 1B-D), coinciding with a marked community differentiation from the undisturbed control reactors (Figs. 3, 4), although only a few taxa are known to participate in autotrophic nitrification. Community evenness, a metric of α- diversity, was suggested to be a key factor in preserving the functional stability of an ecosystem^49^. Therefore, control reactors that displayed better COD removal and complete nitrification with almost no residual 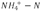 or 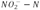, were expected to harbor more diverse communities than treatment reactors. However, we found no significant differences in α-diversity (Fig. 2) in terms of number of ASVs (^0^D) and in how evenly the relative abundances were distributed among taxa in control and treatment reactors (^1^D, ^2^D). The two main genera in both control and treatment reactors (Fig. 3) were *Tetrasphaera* and *Microlunatus,* which include polyphosphate accumulating organisms (PAO) in activated sludge systems. These taxa likely benefited from the high amount of PO_4_^3-^-P in the synthetic wastewater and the alternating anoxic/anaerobic-aerobic cycle conditions in bioreactors. In control reactors, complete nitrification was achieved by the end of an SBR cycle, meaning that after discharging half the reactor volume there was enough nitrate present to support denitrifying organisms like *Paracoccus* and *Alicycliphilus,* which could use it as terminal electron acceptor during the anoxic phase of the cycle. The prevailing nitrifying genera, *Nitrosomonas* and *Nitrospira,* were relatively less abundant in treatment reactors than in control reactors, coincident with a reduction in ammonia and nitrite oxidation activity (details in *Seshan^31^).* On the other hand, organisms like *Candidatus*Competibacter (a glycogen accumulating organism) and *Gemmatimonas* (a PAO) prevailed in treatment reactors, as was also reported for microcosm sludge bioreactors under a similar 3-CA pollutant disturbance^50^. The observed differences in function and dominance of taxa under press disturbed and undisturbed conditions are reasonable, given that organisms face tradeoffs when utilizing resources to maximize their fitness depending on the interactions within their habitat^50,51^.

Community assembly mechanisms were also different between control and treatment reactors, coherent with the idea that underlying assembly mechanisms shape community function and structure^14^. Further, community assembly processes were assessed from ASV data, thus reducing diversity estimation bias by obtaining 10-100 times fewer spurious units than with traditional OTU clustering^34,35^. In our study, disturbance favoured deterministic assembly over time, regardless of the type of null model approach employed for quantification. Phylogenetic-based assembly analysis showed a significant effect of stochastic assembly for control reactors, while treatment reactors displayed dominance of deterministic processes categorized as homogeneous selection^46^ (Fig. 5A). In terms of bacterial turnover assessed via the relative stochastic intensity metric, disturbed reactors showed a stronger role of deterministic mechanisms (Fig. 5B), likely due to the selective pressure via environmental filtering^52^ that occurs under disturbance. Further, β-diversity was always greater than expected for bacterial taxa under the non-phylogenetic null model (Table S2), implying that organisms tended to be more aggregated within replicate bioreactors than expected by chance. Aggregation can be explained by processes of dispersal limitation^53^, due to operating reactors as closed systems without immigration as part of the experimental design of this study, and by habitat filtering^54^, owing to the regular conditions at undisturbed and disturbed levels. Observing similar trends for the effects of disturbance on community assembly via both methods employed is not trivial, as the results from null model assessments are dependent on the algorithms, models and diversity metrics used^55^. The βNTI approach has the advantage of including phylogeny in the analysis^16,46^, but it does not take into account all independent replicates for a given time point to build the null model distribution like the effective bacterial turnover null modelling approach^21,48^. Therefore, the combination of both methodologies allowed us to better exploit the replicated design of this study. Further, differences in community assembly mechanisms operating at phylogenetic and non-phylogenetic levels could be the reason why the acclimation phase showed different trends in assembly mechanisms for each of these methodologies employed.

Our findings concur with those of previous studies of press disturbance in sludge bioreactors using the same micro pollutant (3-CA) at different spatial and temporal scales, as well as different methodologies to assess community structure and assembly. A mesocosm study using 2-L membrane bioreactors, operated during 70 days following a 6-month acclimation period, reported higher similarity among disturbed reactors than control reactors as measured by T-RFLP analysis of a 16S rRNA gene amplicon^11^, although without quantifying assembly mechanisms. Observations derived from community dissimilarities (*i.e.*, patterns of β-diversity) can highlight deterministic effects but have low power to infer stochasticity^23^, the reason why it is important to partition community assembly mechanisms via null model analyses. A recent study quantified assembly mechanisms via null model analysis on metagenomics genus-level data^21^, finding higher stochastic intensity for sludge microcosm reactors that were undisturbed compared to those that were press disturbed with 3-CA. The authors employed a bioreactor scale (20 mL) two orders of magnitude smaller than this work (4 L) and employed a shorter experimental period (35 d) without acclimation. Diversity is multidimensional and scale-dependent^56^, thus assembly mechanisms and the results of null model analyses can differ at different spatial and temporal scales^20,55^. Hence, the observed similar effects of press disturbance in community assembly found at different temporal and volume (*i.e.*, microcosm and mesocosm) scales are relevant. Disturbance was also found to promote deterministic mechanisms in a mesocosm study that used a similar replicated bioreactor system of 5-L SBRs operated for 127 d (including 53 d of acclimation) and the same combination of null-model approaches to quantify assembly mechanisms, but employed a change in organic loading as press disturbance^17^, instead of a xenobiotic. Our findings here strengthen the utility of exploring the role of disturbance in stochastic and deterministic assembly mechanisms for microbial communities.

Overall, this study advances our understanding of the response of complex microbial systems to a xenobiotic compound, by using a joint evaluation of assembly mechanisms, community structure and function of bacterial taxa. In our replicated mesocosm bioreactor system, disturbance altered community function and structure and elicited stronger deterministic mechanisms of community assembly as assessed by two different null model approaches. While this study employed the addition of an aromatic pollutant in the bioreactor feed as a type of press disturbance, more research is needed on different types of disturbances *(e.g.,* pH changes, invading taxa and temperature variations) within different complex microbial systems to further test the general validity of our observations. In this manner, studies covering different temporal and spatial scales, environments and types of disturbance could lead to a general understanding of how press disturbances shift the function, structure, and assembly mechanisms of microbial communities.

## Acknowledgements

This research was supported by the Singapore National Research Foundation and Ministry of Education under the Research Centre of Excellence Program. We thank DI Drautz-Moses for her support with the 16S rRNA gene amplicon library preparation and sequencing pipelines employed.

## Author Contributions

HS designed the experiment and SW obtained the funding for the study. HS performed the experiments. HS and ES performed the molecular work in preparation for amplicon sequencing. ES did the bioinformatics and null model analyses. ES interpreted the data and elaborated the main arguments in the manuscript. All authors contributed to manuscript writing and editing.

## Data availability

DNA sequencing data will be available from NCBI BioProjects upon acceptance of this manuscript. See supplementary information for additional information on null model and multivariate analyses, and data rarefaction.

## Competing interests

The authors declare no competing interests.

## Notes

### Competing Interest Statement

The authors have declared no competing interest.

