## supplementary information for "Press xenobiotic disturbance favors deterministic assembly with a shift in function and structure of bacterial communities in sludge bioreactors"

**Affiliations:**

[†]Current affiliation: Stantec Australia Pty Ltd, 52 Merivale St, South Brisbane QLD 4101, Australia.

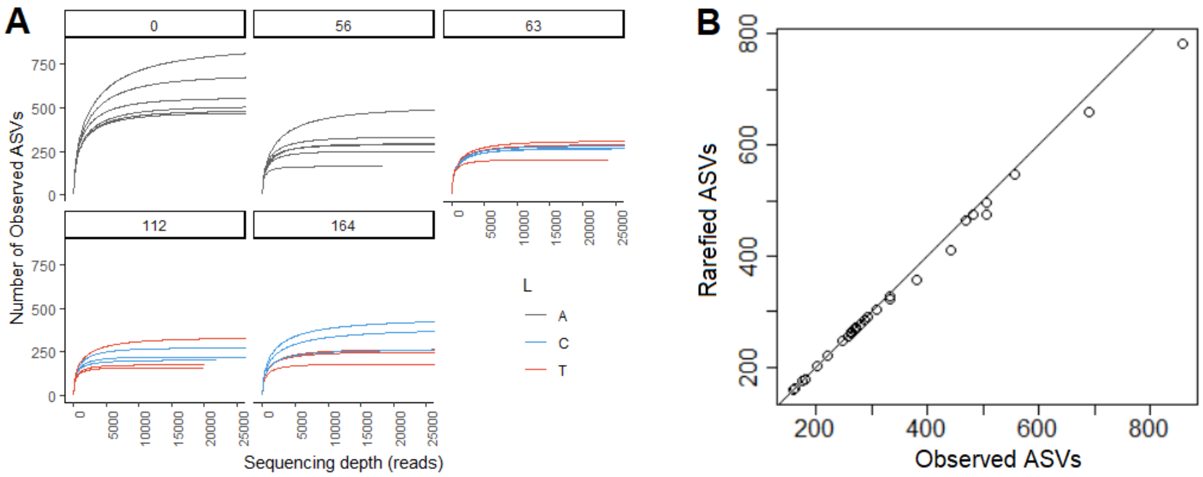


**Fig. S1** – Rarefaction plots for 16S rRNA gene data. (**A**) Rarefaction curves for ASVs separated by time-point sampled. Reactor type: A, acclimation (grey, n = 6); C, control (blue, n = 3); T, treatment (red, n = 3). (**B**) Rarefied versus observed number of ASVs.

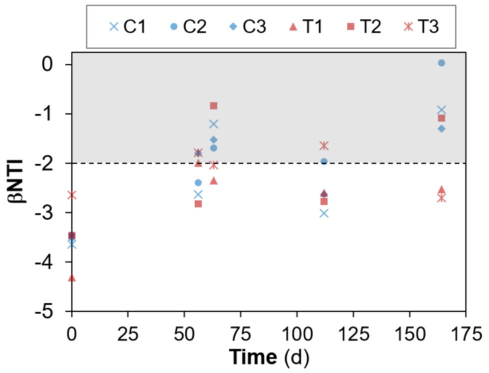


**Figure S2**. Nearest taxon index (βNTI) temporal dynamics for bacterial ASVs, derived from null model analysis. Reactor type: A, acclimation (grey, n = 6); C, control (blue, n = 3); T, treatment (red, n = 3). Symbols represent different independent replicate reactors. Zones where stochastic processes dominate (|βNTI| < 2) are shaded in grey.

**Table S1.** Multivariate tests for relative abundances of bacterial communities (ASV level)[Ψ]

| **Time point** | **No of levels*** | ***n*[‡]** | **df[§] res** | **PERMANOVA[†]** | | **PERMDISP[†]** | |
|---|---|---|---|---|---|---|---|
| | | | | **F** | **P (MC)[¶]** | **F** | **P (perm)** |
| d63 | 2 | 3 | 4 | 0.66 | 0.638 | 0.12 | 0.80 |
| d112 | 2 | 3 | 4 | 5.27 | 0.016[‖] | 0.94 | 0.41 |
| d164 | 2 | 3 | 4 | 5.70 | 0.015[‖] | 0.02 | 1 |

[Ψ]The BC dissimilarity metric was used on square-root transformed data.

* Factor levels: Control reactors (C) and treatment reactors (T).

[†] Number of permutations used was 9,999

[‡] Number of replicates per level

[§] Degrees of freedom of the residual

[¶] Approximate P-value from Monte Carlo permutations

[‖] Significant P-values after correction for multiple comparisons at a False Discovery Rate of 5%, using Benjamini-Hochberg's method.

**Table S2.** Parameter output from non-phylogenetic null model analysis to quantify stochastic intensity

| P[†] | $n$[‡] | d[§] | Bacterial ASVs[¶] | | | | | |
|---|---|---|---|---|---|---|---|---|
| | | | $\boldsymbol{\gamma_{obs}}$ | $\overline{\boldsymbol{\alpha_{obs}}}$ | $\boldsymbol{\beta_{obs}}$ | $\overline{\boldsymbol{\beta_{exp}}}$ | **SI** (%) | $\overline{\boldsymbol{\beta_{exp}}}$:$\boldsymbol{\beta_{obs}}$ |
| A | 3 | 0 | 824 | 511 | 0.3794 | 0.1190 | 31.4 | 0.31 |
| | | 0 | 1068 | 662 | 0.3798 | 0.1640 | 43.2 | 0.43 |
| | | 56 | 597 | 316 | 0.4701 | 0.1409 | 30.0 | 0.30 |
| | | 56 | 495 | 292 | 0.4108 | 0.0753 | 18.3 | 0.18 |
| C | 3 | 63 | 459 | 272 | 0.4067 | 0.0836 | 20.6 | 0.21 |
| | | 112 | 410 | 232 | 0.4350 | 0.0523 | 12.0 | 0.12 |
| | | 164 | 652 | 353 | 0.4591 | 0.1596 | 34.8 | 0.35 |
| T | 3 | 63 | 465 | 266 | 0.4280 | 0.0843 | 19.7 | 0.20 |
| | | 112 | 398 | 221 | 0.4439 | 0.0796 | 17.9 | 0.18 |
| | | 164 | 418 | 231 | 0.4482 | 0.0798 | 17.8 | 0.18 |

[†] Reactor type: A, acclimation; C, control; T, treatment.

[‡] Number of independent replicates

[§] Time (days)

[¶] Null model parameters: $\boldsymbol{\gamma_{obs}}$, observed gamma diversity; $\overline{\boldsymbol{\alpha_{obs}}}$, mean observed alpha diversity; $\boldsymbol{\beta_{obs}}$, observed beta diversity; $\boldsymbol{\beta_{obs}}$, observed beta diversity; SI, stochastic intensity; $\overline{\boldsymbol{\beta_{exp}}}$:$\boldsymbol{\beta_{obs}}$, expected (mean) to observed beta diversity ratio.
